# Bioinformatics analysis of Uncyclotides: a potential antimicrobial peptide to protect against heart failure diseases

**DOI:** 10.1101/2023.10.23.563516

**Authors:** Carlos Eliel Maya-Ramírez, Muhammad Sufyan, Abdullah R Alanzi

## Abstract

Heart failure (also known as HF) is a clinical disorder triggered by functional and structural defects in cardiac muscles causing blood ejection or damage of ventricular filling. Even though the progression in the field of medicine, HF control, which generally presents as a syndrome, has been a task to healthcare sources. This is reproduced by the comparatively higher ration of readmissions with enhanced disease and death linked with HF. Congestive heart failure is also linked with expressional changes of human uncoupling protein proteins. So, we need to tackle the heart failure problem with the help of bioinformatics analysis. In this study, we retrieved three antimicrobial peptides (Cathelicidins, CRAMP and Uncyclotides) and four human uncoupling proteins (UCP 1, UCP 2, UCP 3 & UCP 4) from DRAMP and NCBI web server. Physiochemical properties and domains analysis of UCPs and antimicrobial peptides was performed by using Protparam, DRAMP and InterPro databases correspondingly. To predict the UCPs structure, homology modelling was performed by SWISS-MODEL server and structural assessment was done by Ramachandran plots. We detected the best antimicrobial peptide (Uncyclotides) against heart failure diseases after HPEPDOCK analysis. We also confirmed stability of UCP 1-Uncyclotides complex, and shown the slight distortion of the deformability plot supported by highest eigenvalues after Molecular Dynamics analysis. Therefore, this in silico approach identifies potential antimicrobial peptides “Uncyclotides” (acts as inhibitors) of heart failure diseases, which have to be confirmed further after experimental and clinical trials.

## Introduction

The considerable pervasiveness and substantial difficulties of heart failure have caused a simple and translational study regarding the disease etiology and therapeutic choices for cure of heart failure diseases (Benjamin et Schneider, 2005) . Various studies have revealed that the heart failure progress and its evolution are linked with expressional changes of numerous genes, comprising metabolic, signaling and structural genes (Hwang et al., 2002). Generally, it is not recognized whether these changes in gene expression (Chen et al., 2003), usually from adult to fetal phenotypes (Depre et al., 1998), are useful or harmful to the heart failure (deGoma et al., 2006).

Family of uncoupling proteins (also known as UCPs) belongs to specific proteins that are associated with heart failure diseases especially when UCPs shown down regulation (UCP 2-3). These proteins involved in the production of ROS (known as Reactive oxygen species). Consequently, UCPs could possibly play a significant part in heart failure disease, because of the factuality that the failing heart is linked with two main anomalies such as process of energy conversion and enriched oxidative stress. Both anomalies may cause apoptosis, an emblem characteristic of heart failure at cellular level (Laskowski et Russell, 2008).

The foremost described UCP 1 (may be known as thermogenin), was found to disperse the gradient of protons in an “energy wasting” way, producing heat in its place of ATP. Mainly limited to the brown adipose tissue, UCP 1 is involved in the process of non-shivering thermogenesis. Afterwards, numerous proteins same to UCP 1 were revealed, forming a UCPs family. The UCP 2 was recognized as a UCP 1 structural homolog (Gimeno et al., 1997). Throughout the human body, UCP 2 is present in many tissues, also in heart (depends on species). Similar to UCP 2 (Fleury et al., 1997), UCP3 is present in inner side of mitochondrial membrane with substantial structural homology to UCP1 and also remains in heart muscle. On protein level, UCP3 is > 70% similar to UCP2 and > 50% similar to UCP1 (Vidal-Puig et al., 1997). In advanced states, the utmost frequent cause of heart failure is related to ischemic heart ailment. About 50% cases, the origin of heart failure are attributed to preceding myocardial infarction on the basis of population study and also investigation methods (Fox et al., 2001).

Antimicrobial peptides (also known as AMPs or peptide of genetically coded antibiotics), are conserved components associated with innate immunological responses (Zasloff, 2002). The AMPs are formed in several living being like as amphibians, animals, plants, diverse bacteria’s, insects and also in vertebrates, where they depict an immunological defense system (Brogden, 2005). In vertebrates, these AMPs perform different functions especially in immune system (few phagocytic cells destroy or engulf invasive bacteria’s and also stop/avoid the settlement of host tissues) (Ganz, 1999). From the last few decades, many families of AMPs produced from diverse living organisms have been reported. But in human beings, three main diverse groups were described like as cathelicidins, defensins and also histatins. Further antimicrobial peptides (dermacidin, granulysin, lactoferrin and so on) have been recognized. These peptides perform effective functions against huge number of diverse microorganisms (diverse bacteria’s, viruses, fungi) (Boman, 2000). Several antimicrobial peptides or proteins have been linked with evolving global issues (diabetes, heart diseases and also obesity) (Zaiou, 2007).

Development of congestive heart failure (also known as CHF) is linked with expressional changes of an extensive range of diverse structural, metabolic and signaling proteins. Such a one consequence is the lower expression level of UCPs in the heart failure site. This proteins group controls the potential of mitochondrial membrane and thus performs a main part in process of energy metabolism and also production of ROS by mitochondria (Laskowski et al., 2008). That’s why, we deciding to do work on UCPs. The main objective of this study is to identify the potential antimicrobial peptide to target heart failure disease. For bioinformatics analysis, we will use three antimicrobial peptides (Cathelicidin, CRAMP and Uncyclotides) and four human uncoupling proteins (UCP 1, UCP 2, UCP 3 & UCP 4). For structure prediction, molecular docking and dynamics analysis, we will explore diverse bioinformatics software’s/webserver/tools and will select the best antimicrobial peptide and will be used in pharmaceutical industries to tackle the issue of heart failure.

## Material and Methods

### Data retrieval

We retrieved antimicrobial peptides associated with cardiovascular diseases like as Cathelicidins (DRAMP18429), CRAMP (DRAMP03404) and Uncyclotides (DRAMP00926) from DRAMP web server (http://dramp.cpu-bioinfor.org/). The DRAMP server provides information about AMPs/peptides name, sequences, source and also their activity. We also retrieved four human uncoupling proteins associated with heart disease (UCP1 (NP_068605.1), UCP2 (P55851.1), UCP3 (NP_003347.1) & UCP4 (NP_001190981.1)) from NCBI (https://www.ncbi.nlm.nih.gov/protein/).

### Analysis of Physiochemical properties

Two different online web servers (ProtParam and DRAMP web servers) were used to assess the physiochemical properties of human UCPs and antimicrobial peptides. To access the both severs, For UCPs and antimicrobial peptides; we used both ProtParam (https://web.expasy.org/protparam/) and DRAMP (http://dramp.cpu-bioinfor.org/) web servers respectively.

### Domain analysis

For prediction of human UCPs domains, we used InterPro database (https://www.ebi.ac.uk/interpro/). This database provides information regarding proteins functional analysis (also known as PFA) by grouping them into diverse families and also predicting significant domains and sites.

### Homology modelling

For homology modeling of human uncoupling proteins, we used SWISS-MODEL server (https://swissmodel.expasy.org/). The predict structures of uncoupling proteins was validated by using Ramachandran plots. This server used both supervised machine learning and Support Vector Machines algorithms for comparative modelling.

### Molecular docking analysis using MDockPeP server

Molecular docking analysis of four human uncoupling proteins (UCP1, UCP2, UCP3 & UCP4) with antimicrobial peptides (Cathelicidins, CRAMP & Uncyclotides) by using “HPEPDOCK” web server (http://huanglab.phys.hust.edu.cn/hpepdock/) and then selected best combination of docking complex with the help of docking energy scores.

### Molecular dynamics analysis

After docking, we performed molecular dynamics analysis of best docked complex using iMODS web server (https://imods.iqfr.csic.es/). This server does normal mode analysis on internal coordinates of biological molecules like as DNA and protein structures.

## Results and discussion

### Physiochemical properties of UCPs and antimicrobial peptides

Physiochemical properties of human UCPs and antimicrobial peptides were calculated by using ProtParam (https://web.expasy.org/protparam/) and DRAMP tool (http://dramp.cpu-bioinfor.org/) respectively. These properties include aliphatic index, isoelectric point, length, molecular weight, molecular formula GRAVY, Instability index, estimated half-life and many more. According to ProtParam results, UCP 1, UCP 2 and UCP 4 were stable but UCP 3 was unstable protein. The estimated half-life of all UCPs was same (30 hours in mammalian reticulocytes, >20 hours in yeast & >10 hours in E.coli) (Table 1).

**Table 1:** Physiochemical properties of human Uncoupling proteins.

| Parameters | UCP 1 | UCP 2 | UCP 3 | UCP 4 |
| --- | --- | --- | --- | --- |
| Length | 307aa | 309 aa | 312aa | 245aa |
| Atomic composition | C= 1477, H=2382, N=394, O=427 & S=16 | C= 1476, H=2353, N=413, O=427 & S=16 | C= 1529, H=2428, N=412, O=435 & S=21 | C= 1220, H= 1938, N=340, O=343 & S=11 |
| Molecular weight | 33004.50da | 33229.39da | 34215.86da | 27209.54da |
| Isoelectric point | 9.26 | 9.74 | 9.31 | 8.61 |
| Aliphatic index | 95.57 | 82.46 | 84.97 | 93.14 |
| GRAVY | 0.206 | 0.067 | 0.075 | -0.120 |
| Molecular formula | C<sub>1477</sub>H<sub>2382</sub>N<sub>394</sub>O<sub>427</sub>S<sub>16</sub> | C<sub>1476</sub>H<sub>2353</sub>N<sub>413</sub>O<sub>427</sub>S<sub>16</sub> | C<sub>1529</sub>H<sub>2428</sub>N<sub>412</sub>O<sub>435</sub>S<sub>21</sub> | C<sub>1220</sub>H<sub>1938</sub>N<sub>340</sub>O<sub>343</sub>S<sub>11</sub> |
| Total number of atoms | 4696 | 4685 | 4825 | 3852 |
| Total number of negatively charged residues | 20 | 19 | 22 | 26 |
| Total number of positively charged residues | 30 | 34 | 33 | 28 |
| Estimated half-life | 30 hours in mammalian reticulocytes, >20 hours in yeast & >10 hours in E.coli | 30 hours in mammalian reticulocytes, >20 hours in yeast & >10 hours in E.coli | 30 hours in mammalian reticulocytes, >20 hours in yeast & >10 hours in E.coli | 30 hours in mammalian reticulocytes, >20 hours in yeast & >10 hours in E.coli |
| Instability index (II) | 37.97 (stable) | 41.95 (unstable) | 39.31 (stable) | 37.97 (stable) |

On the other hand, physiochemical properties of antimicrobial peptides includes DRAMP ID, sequence, length, Biological activity, Mass, PI, Boman index, Hydrophobicity, aliphatic index, half-life and many more (Table 2). All the peptides shown antimicrobial activities but cathelicidins and CRAMP also showed antibacterial and antifungal activities. The half-life of cathelicidins and CRAMP was same (Mammalian: 30hrs, Yeast: >20hrs, E.coli: >10hrs) but little bit different in Uncyclotides (Mammalian: 1.9hrs, Yeast: >20hrs, E.coli: >10hrs).

**Table 2:**
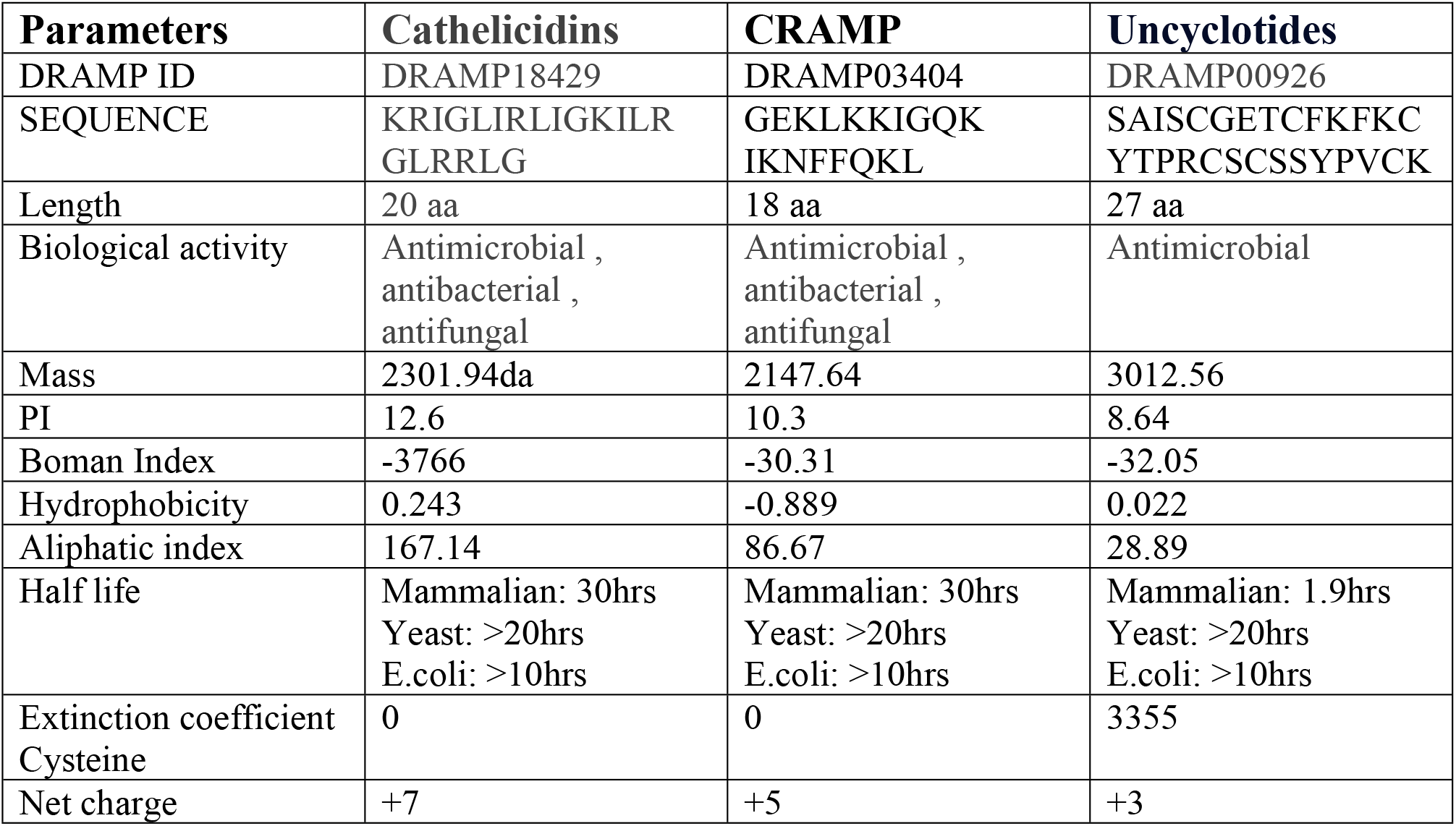
Physiochemical properties of Antimicrobial peptides.

### Domain analysis of UCPs

Domains analysis of human UCPs was performed by using InterPro database (https://www.ebi.ac.uk/interpro/)(Mulder et Apweiler, 2008). Basically domains annotation is a substitute method to search sequence similarity of proteins. Protein domains are diverse regions of their sequences which are extremely preserved through an evolution process. Domains are involved in protein-protein association and also perform critical functions in interaction network (Lee et Lee, 2009). Domains are represented by different colors like as Mit_uncoupling_UCP_like (18-30, 47-66, 100-112, 166-183, 209-227 and 286-307) by yellow, mitochondrial brown fat uncoupling protein signature domains by blue in Family (six domains at diverse positions), but in homologous superfamily, Mt_carrier_dom_sf, mitochondrial carrier, and mitochondrial carrier domain by maroon, light yellow and dark green colors respectively. In repeat, Mitochondrial_sb/sol_carrier (11-109, 114-207, 212-299), solute carrier (Solcar) repeat profile and mitochondrial carrier protein denoted by dark green, light green and light blue colors respectively. In the unintegrated, mitochondrial uncoupling protein 2 and Mitochondrial dicarboxylate carrier-related domains denoted by purple and moderate blur colors respectively. Protein function was also predicted by InterPro GO terms like as Mitochondrial transport as a biological process (GO: 0006839) and mitochondrial membrane as a cellular component (GO: 00031966) (Figure 1).

**Figure 1:**
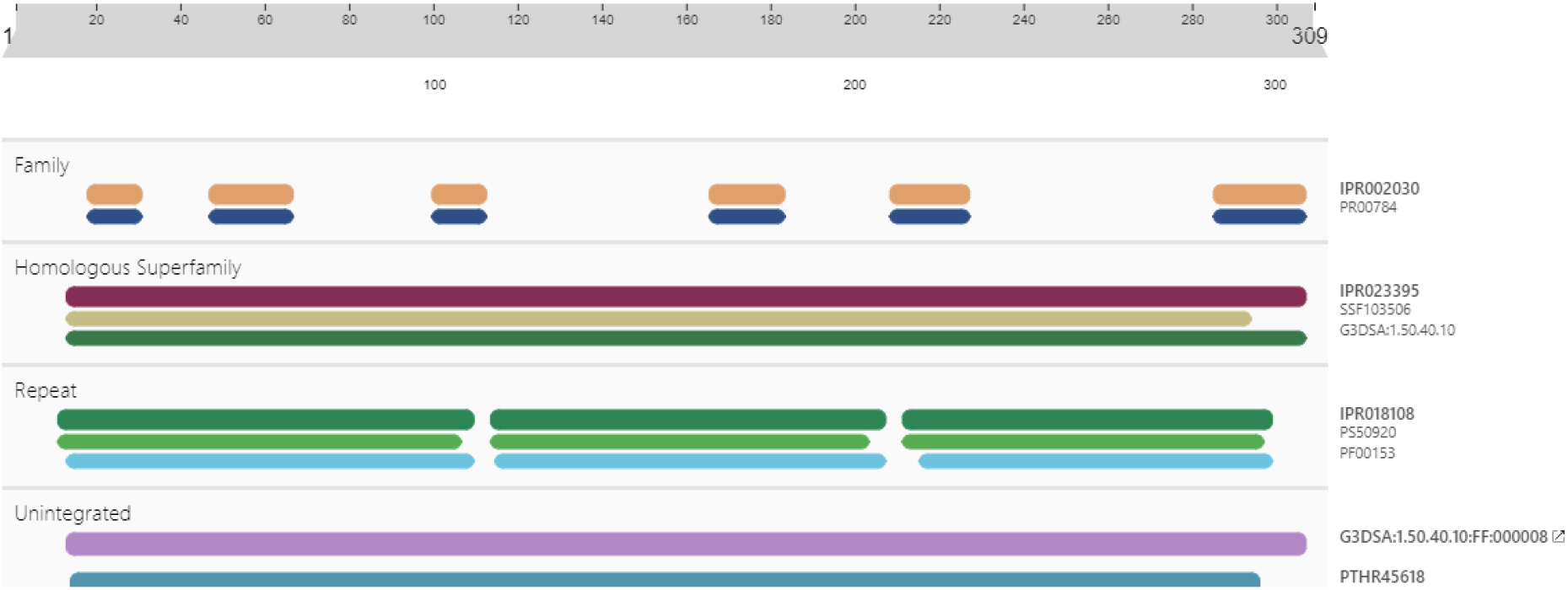
Domain analyses of human UCPs, domains are represented by different colors like as Mit_uncoupling_UCP_like by yellow and mitochondrial brown fat uncoupling protein signature by blue color.

### UCP Structure prediction by Homology modelling

Amino acid sequences of four human uncoupling proteins (UCP 1 -4) retrieved from NCBI **(https://www.ncbi.nlm.nih.gov/protein/)** (The NCBI Accession No: for each protein was NP_068605.1 for UCP 1, P55851.1 for UCP 2, NP_003347.1 for UCP 3 and NP_001190981.1 for UCP 4). For protein comparative modelling, we used four UCPs and predicted their structures by using web server “SWISS-MODEL” (Schwede et al., 2003). This server automatically selects the templates for human uncoupling proteins, generates proteins structures with also providing structure assessments (by making Ramachandran plots with MolProbity scores). Predicted structure and their assessment of human uncoupling proteins were given in Figure 2(A-H). For prediction of 3D models, energy minimization was done to decrease steric crashes and strains without considerably changing the overall protein structure. The calculation of energy and their minimization were performed by using GROMOS96 force field (Scott et al., 1999) and executing Swiss-PDB Viewer. After optimization, predicted structures were verified by plotting Ramachandran plots (Sasisekharan, 1963).

**Figure 2:**
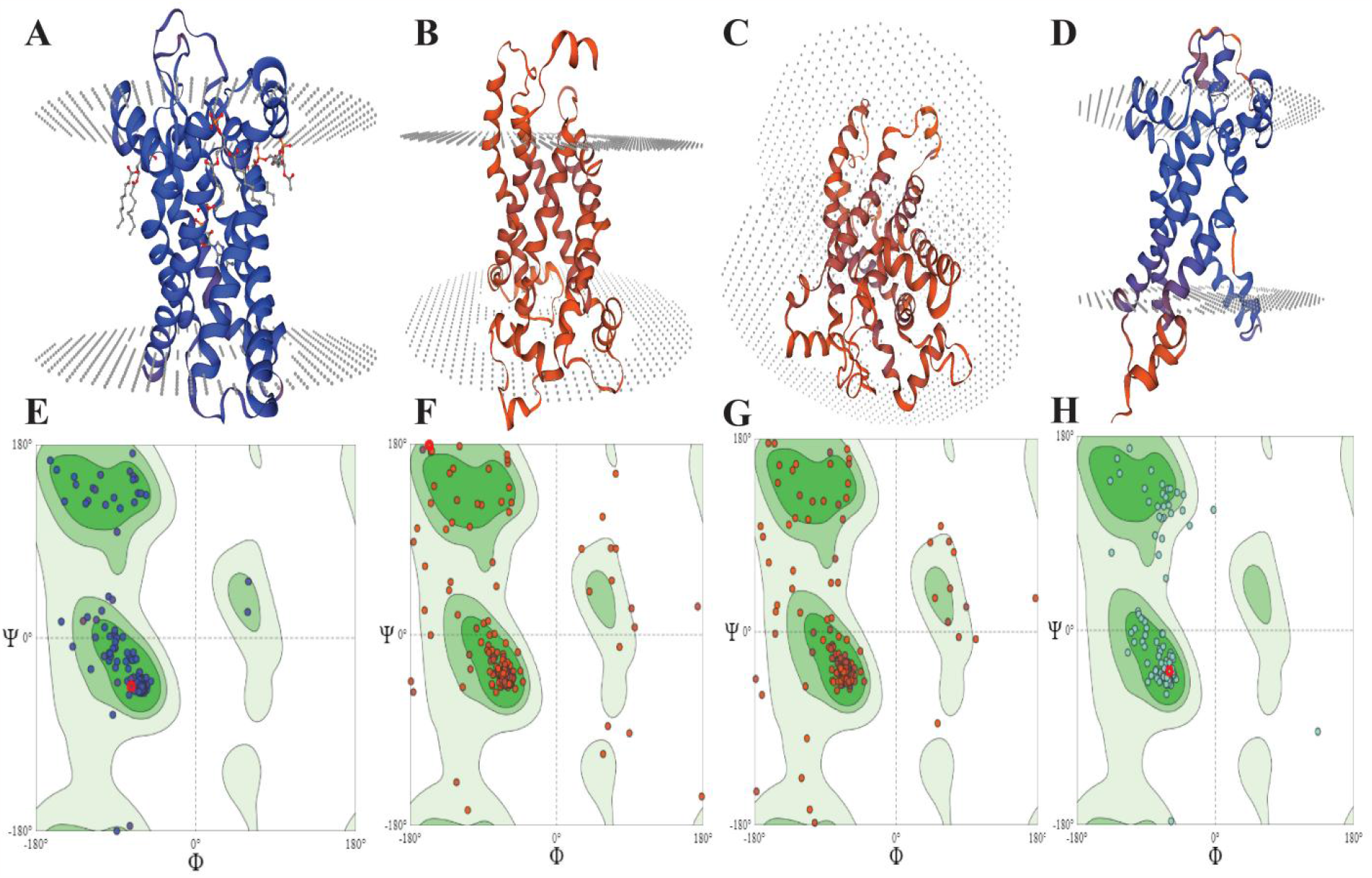
Predicted structures of human uncoupling proteins predicted by using SWISS-MODEL, 2A, represented 3D structure of UCP 1; 2B, 3D structure of UCP 2; 2C, 3D structure of UCP 3; and 2D, 3D structure of UCP 4; 2E, Structural assessment of UCP 1 using Ramachandran plots of UCP 1; 2F, Ramachandran plots of UCP 2; 2G, Ramachandran plots of UCP 3; 2F, Ramachandran plots of UCP 4.

**Figure 3:**
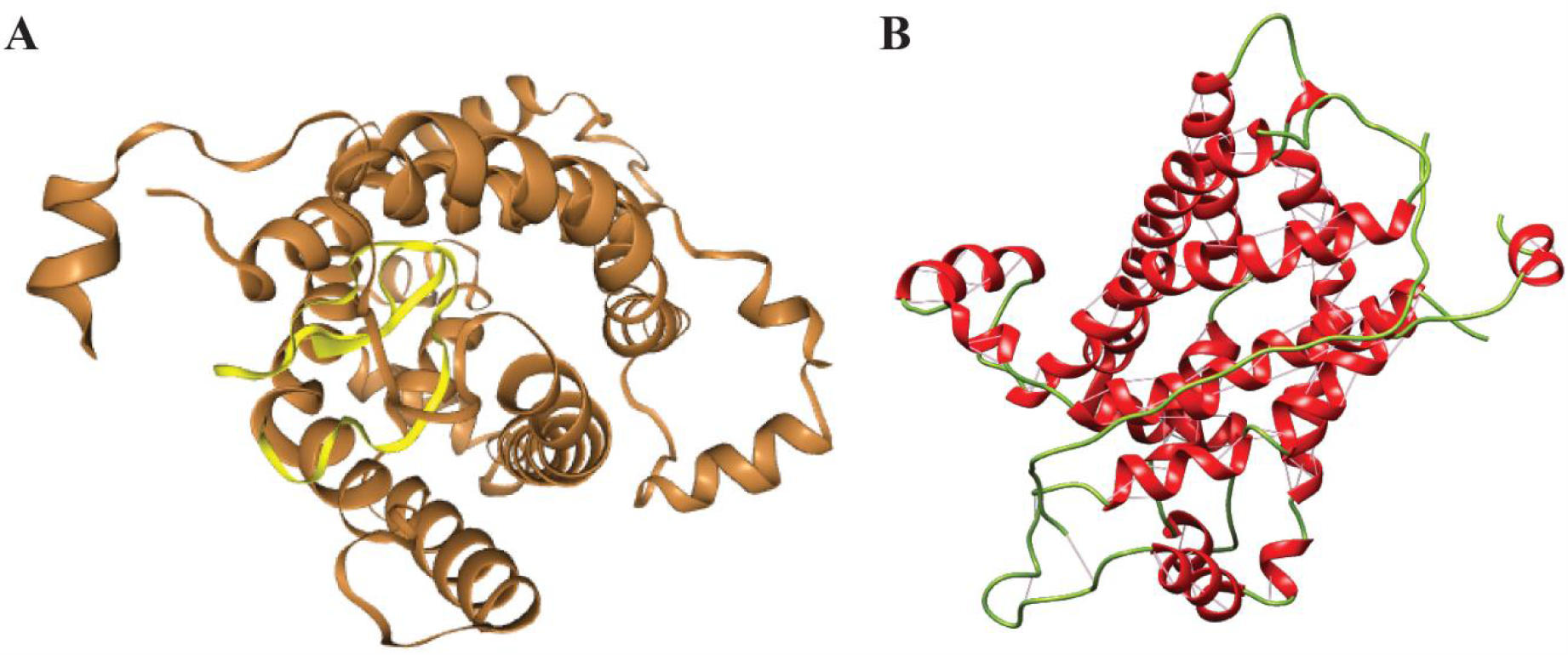
4A, Best docking complex (UCP 3-Uncyclotides) by HPEPDOCK, UCP 3 protein and Uncyclotides represented by brown and by yellow color respectively; 4B, representing 2D image of docking complex, in which Helix, Strand and coil shown by red, blue and green color.

In Ramachandran plots (Figure 2 E-H), 98.26% residues are present in favorable regions of UCP1 with 1.04 MolProbity score. The UCP 2 has 88.05% residues in favorable regions (with 2.13 MolProbity score), UCP 3 has 87.88 % residues in favorable region (with 2.23 MolProbity score) and also UCP 4 has 94.96% residues in favorable regions (with 1.11 MolProbity score). There is not any bad bonds are reported in UCP3 and UCP4. On the other hand, UCP1 has two (A296 LEU, A212 VAL) with zero C-Beta Deviations and UCP 2 has three bad bonds (A48 GLN-A49 GLY, A48 GLN) with three C-Beta Deviations in their predicted structure (Table 3).

**Table 3:** Prediction of Ramachandran parameters.

| **Parameters** | **UCP 1** | **UCP 2** | **UCP 3** | **UCP 4** |
| --- | --- | --- | --- | --- |
| MolProbity Score | 1.04 | 2.13 | 2.23 | 1.11 |
| Clash Score | 2.55 | 0.67 | 1.94 | 0.80 |
| Ramachandran Favoured | 98.26% | 88.05% | 87.88% | 94.96% |
| Ramachandran Outliers | 0.00% | 4.44% | 4.71% | 2.52% |
| Rotamer Outliers | 0.41% | 11.69% | 7.63% | 1.03% |
| C-Beta Deviations | 0 | 3 | 4 | 1 |
| Bad Bonds | 2 | 3 | 0 | 0 |

### Protein-peptide docking by using HPEPDOCK 2.0 web server

Molecular interaction of protein-peptide performs very critical part in numerous cellular functions. Thus, structure prediction of docking complexes is very important for understanding the diverse molecular functions with related biological methods and also in producing peptide drugs. The HPEPDOCK2.0 web server used hierarchal algorithm for blind protein-peptide docking (P. Zhou et al., 2018). This web server used for protein-peptide blind docking on the basis of hierarchical algorithm. By using HPEPDOCK scores, Peptide-1 (have -241.390 docking score) and Peptide-2 (having -241.619 docking score) have highest docking score with respect to other given peptides, whereas ACE2 (-351.74) has the uppermost docking energy score with the RBD compound (Mukhopadhyay et Sarkar, 2021).

Successively, the top ten docking models are mostly reflected the most substantial predictions of docking results; the results comprises an interactive vision of the top docking models by Jmol software (Hanson et al., 2013). Lower the docking energy score, higher the binding affinity of UCPs with antimicrobial peptides. According to the docking energy scores calculated by HPEPDOCK 2.0 web server, we conclude that UCP 3 showing best interaction results with “Uncyclotides” peptide (-256.739Aº) (Figure 4 & Table 4). After UCP3-Uncyclotides, both UCP 4-Uncyclotides (-250.883Aº) and UCP4-Cathelicidins (-248.656Aº) were shown better interactions.

**Table 4:** Docking results calculated by HPEPDOCK.

| Sr. no | UCPs Protein- antimicrobial peptides | Docking scores |
| --- | --- | --- |
| 1 | UCP1-Cathelicidins | -222.486 |
| 2 | UCP1-CRAMP | -189.305 |
| 3 | UCP1-Uncyclotides | -229.965 |
| 4 | UCP2-Cathelicidins | -242.270 |
| 5 | UCP2-CRAMP | -195.439 |
| 6 | UCP2-Uncyclotides | -230.182 |
| 7 | UCP3-Cathelicidins | -191.956 |
| 8 | UCP3-CRAMP | -178.262 |
| **9** | **UCP3-Uncyclotides** | **-256.739** |
| 10 | UCP4-Cathelicidins | -248.656 |
| 11 | UCP4-CRAMP | -183.594 |
| 12 | UCP4-Uncyclotides | -250.883 |

**Figure 4:**
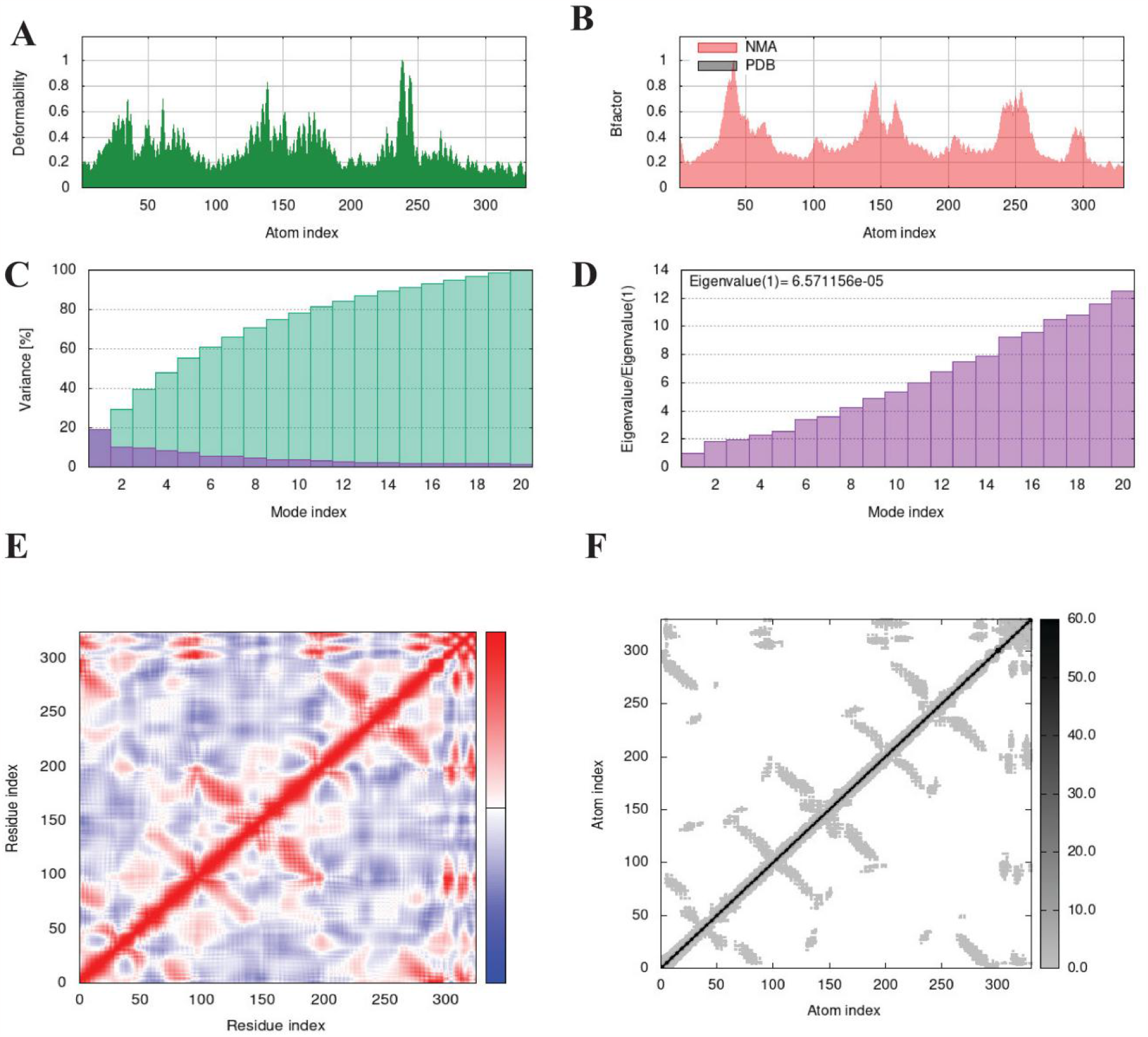
Molecular dynamics analysis of UCP3-Uncyclotides using iMODS server. (A) Deformability plot of UCP3-Uncyclotides. (B) B-factor plot. (C) Eigenvalue plot of UCP3-Uncyclotides. (D) Variance plot. (E) Covariance matrix plot in which correlation, anti-correlation and un-correlation of residual motion represented by red, blue and white color correspondingly. (F) Elastic network model of UCP3-Uncyclotides, more rigid spring and soft spring represented by dark grey and white color respectively.

### Molecular dynamics analysis of UCP 3-Uncyclotides docked complex

Molecular dynamics analysis by iMODS server was used to assess the firmness and motion of UCP 3-Uncyclotides docking complex. In Figure 5A, deformability plot indicates that a slight distortion was reported in the UCP 3-Uncyclotides, as showed by hinges. This plot complemented by the B-factor/motility plot, which is proportional to the root mean square (also known as RMS) and displays the firmness of the UCP 3-Uncyclotides docked complex (Figure 5B). Moreover, a higher eigenvalue of 6.571156×10−5 was detected, which displays the energy requirement to distort the UCP 3-Uncyclotides docked complex, and also proposes complex stability (Figure 5C) and variance graph (Figure 5D). The graph of covariance matrix among the residues pairs is shown in Figure 5E; in which red, blue and white color denote correlation, anti-correlation, and un-correlation motion of residues, correspondingly. The elastic network model of the UCP 3-Uncyclotides docked complex presents the complex stiffness. Higher protein stiffness represented by darker gray in specific regions and lower protein stiffness by white color (Figure 5F). From MDA results, we proved the stability of UCP 3-Uncyclotides complex, and also shown by the slight distortion of the deformability plot supported by highest eigenvalues.

## Discussion

In this work, we have described diverse protein domains and its structural information, homology modelling, molecular docking and dynamics analysis of the four human uncoupling proteins (UCP 1, UCP 2, UCP 3 & UCP4) and three peptides (Cathelicidins, CRAMP and Uncyclotides used as ligands) and we have revealed how that specific information can be utilised *in silico* to identify prospective therapeutic drug targets for heart failure ailments. During this research work, we have reflected and also justified the information extracting from both proteins domains and complete domain structure. Different databases related to protein domains and family has preserved a critical site in the ecology of in silico biological databases/servers/tools and other resources (Mulder et al., 2003). So, we performed domain analysis on human uncoupling protein using InterPro database (Apweiler et al., 2001) and determined that the UCPs (Teshima et al., 1999) belonged to Mit_uncoupling_UCP_like domains, mitochondrial brown fat uncoupling protein signature domains (Family), and Mt_carrier_dom_sf (homologous superfamily).

Molecular docking is an effective tool extensively utilised to see the molecular features of docked complex interactions (Kitchen et al., 2004). This docking is used to assess the different peptides inside the active sites of proteins such that the receptor proteins remain constant and maintain their unique structure (Sharfalddin et al., 2021). As well as, our work displays that the three antimicrobial peptides; Cathelicidins, CRAMP and Uncyclotides; and the Uncyclotides interact effectively with the UCP 1 and UCP 3 and UCP 4 receptor proteins, but Uncyclotides showing best results with UCP 3 protein and also revealing diverse therapeutic modes of action through potentially regulating theses receptors.

All the antimicrobial peptides, except CRAMP, have more interactions with UCP 3 and UCP 4 than the UCP 1. Uncyclotides displayed the best docking scores succeeded by Cathelicidins and CRAMP. The evaluation of molecular docking to UCP 3 and UCP 4 has unlocked a unique feature in this respect and its outcomes should be confirmed through an investigational study. Downregulation of human uncoupling proteins are associated with heart failure disease (Laskowski et al., 2008). So, after docking analysis, we predicted that Uncyclotides will be bound to the human uncoupling proteins and also enhance the expression of UCPs and helps to prevent heart failure diseases in future. Diverse significant residues on the basis of structures and functions like as binding residues of inhibitor at the best site of docked complex (CXCR4-complex & CXCR7-complex) were recognized (Murad et al., 2022). In recent times, it was described that activation of CXCR7 may be a promising curative target in ischemic myocardium ailment. It secured diverse ischemic cells in hypoxic endothelial cells and infarction model by activating the formation of new blood vessels and also reducing the cell death in C57BL/6 J mouse model of severe heart attack (Zhang et al., 2020). Usually, based upon the protein docking scores it can be determined which peptide/compound, as opposed to others, is very efficient to the preferred receptor or proteins. The compounds/peptides permanency inner side of the cavity and time measurement of docking complex can be assessed by utilising the molecular dynamics simulation, which activates the driving behavior of biological systems with respect to time period (H. Zhou et al., 2011), but unluckily, we do not have the databases which can prepare such work as well as being inefficient since it takes overmuch time to assess the constancy of compounds/peptides within the cavity to 500 ns or maybe not. We also affirmed the strength of docking complex (UCP 3-Uncyclotides), and revealed the slight distortion of the deformability plot linked with highest eigenvalues after molecular dynamic analysis. Therefore, this in silico approach identifies “Uncyclotides” as potential antimicrobial peptides against heart failure ailments, which have to be verified further by clinically and experimentally.

## Conclusion

In this research, Bioinformatics analysis was performed on human uncoupling proteins because expression levels of these proteins linked with congestive heart failure diseases. Physiochemical properties and domains analysis was perform to check the important specification of therapeutic proteins to be established, support drug development process and also provide insight about functional and structural information of uncoupling proteins. Protein Homology Modeling was performed to predict the structures of human uncoupling proteins (UCP 1, UCP 2, UCP 3 & UCP 4) and structures assessment was done by Ramachandran plots. We also screened out the best antimicrobial peptide (Uncyclotides) against heart failure diseases after HPEPDOCK analysis. We also confirmed stability of UCP 3-Uncyclotides complex, and shown the slight distortion of the deformability plot supported by highest eigenvalues after MDS analysis. Therefore, this in silico approach identifies potential antimicrobial peptides “Uncyclotides” of heart failure diseases, which have to be confirmed further (like as experimentally and clinically).

## Acknowledgment

The present study was supported by the Researchers Supporting Project (grant no. RSPD2023R885), King Saud University, Riyadh, Saudi Arabia.

